# Mobile element integration reveals a chromosome dimer resolution system in Legionellales

**DOI:** 10.1101/2021.09.12.459815

**Authors:** Beth Nicholson, Shayna R. Deecker, Alexander W. Ensminger

## Abstract

In bacteria, the mechanisms used to repair DNA lesions during genome replication include homologous recombination between sister chromosomes. This can lead to the formation of chromosome dimers if an odd number of crossover events occurs. The concatenated DNA must be resolved before cell separation to ensure genomic stability and cell viability. The broadly conserved *dif*/Xer system counteracts the formation of dimers by catalyzing one additional crossover event immediately prior to cell separation. While *dif*/Xer systems have been characterized or predicted in the vast majority of proteobacteria, no homologs to *dif* or *xer* have been identified in the order Legionellales. Here we report the discovery of a distinct single-recombinase *dif*/Xer system in the intracellular pathogen *Legionella pneumophila*. The *dif* site was uncovered by our analysis of *Legionella* mobile element-1 (LME-1), which harbors a *dif* site-mimic and integrates into the *L. pneumophila* genome via site-specific recombination. We demonstrate that *lpg1867* (herein named *xerL*) encodes a tyrosine recombinase that is necessary and sufficient for catalyzing recombination at the *dif* site, and that deletion of *dif* or *xerL* causes filamentation along with extracellular and intracellular growth defects. We show that the *dif/*XerL system is present throughout Legionellales and that *Coxiella burnetii* XerL and its cognate *dif* site can functionally substitute for the native system in *L. pneumophila*. Lastly, we describe an unexpected link between *C. burnetii dif*/Xer and the maintenance of its virulence plasmids.

**Significance Statement:** The maintenance of circular chromosomes depends on the ability to resolve aberrant chromosome dimers as they form. In most proteobacteria, broadly conserved Xer recombinases catalyze single crossovers at short, species-specific *dif* sites located near the replication terminus. Chromosomal dimerization leads to the formation of two copies of *dif*, leading to rapid site-specific recombination and elimination of the duplicated intervening sequence. The apparent absence of chromosome-dimer resolution mechanisms in Legionellales has been a mystery to date. By studying a phage-like mobile genetic element, LME-1, we have identified a previously unknown single-recombinase *dif*/Xer system that is not only widespread across Legionellales but whose activity is linked to virulence in two important human pathogens.

## INTRODUCTION

The circular nature of bacterial chromosomes brings about a problem for the partitioning of genetic information to daughter cells: the formation of chromosome dimers during replication. Dimers are generated when homologous recombination-mediated repair of DNA lesions results in an odd number of crossover events between sister chromosomes. The concatenated DNA must be resolved into monomers for proper cell division to occur. The essential and broadly conserved bacterial *dif*/Xer system overcomes this topological problem by catalyzing one additional crossover event immediately prior to cell separation (1, 2). The *dif* (<u>d</u>eletion-induced <u>f</u>ilamentation) site is a conserved ∼30-bp DNA element on the chromosome that contains binding sites for either two copies of a single tyrosine recombinase or one copy of two different ones (3). In a highly regulated process, a nucleoprotein complex containing the Xer tetramer and two aligned *dif* sites forms a recombination synapse, and site-specific recombination between them deconcatenates the chromosome dimer into monomers (4–6).

Dimer resolution via site-specific recombination by tyrosine recombinases was first discovered in plasmids (7, 8) including ColE1, in which a DNA sequence named *cer* was found to be essential for plasmid monomerization and stability (8). Recombination between *cer* sites was later found to require catalysis by XerC (9) and XerD (10) encoded on the *E. coli* genome. These findings, along with homology between the chromosomal *dif* site and *cer*, led to the discovery of the *dif*/XerCD chromosome dimer resolution (CDR) pathway in *E. coli* (10–13) and later in several other species of bacteria (14–19) and archaea (20, 21). Subsequent dissection of the *dif*/Xer recombination machinery has been aided by another group of mobile genetic elements, named integrative mobile elements exploiting Xer (IMEX) (22–27). IMEXs, which include lysogenic phages, genomic islands, and plasmids, contain a *dif*-mimic sequence which allows them to integrate into the chromosome at the *dif* site using the host’s Xer recombination machinery (5, 28).

Homologs of *xerC* and *xerD* have been predicted to be present in the vast majority of proteobacterial species (29). Exceptions to this include two groups of bacteria that utilize a single-recombinase system. The *Streptococci*/*Lactococci* use a single recombinase XerS and an atypical *dif* site (16) and a group of epsilon-proteobacteria, including *Helicobacter* and *Campylobacter* species, uses a single-recombinase XerH system (14, 17, 29). While *dif*/Xer homologs have been detected in almost 90% of proteobacteria, no *dif* or Xer homologs could be detected in the order Legionellales, despite encoding RecA, RecBCD, and RecF, which are thought to be responsible for dimer formation during replication (29). How then do the members of the order Legionellales overcome the threat to genome stability that chromosome dimers impose? It is possible that they use an entirely different method for CDR, or that its *dif*/Xer system is sufficiently divergent from others to avoid detection by sequence-based homology searches.

Several strains of *Legionella pneumophila* harbor a phage-like integrative mobile genetic element, named *Legionella* mobile element-1 (LME-1) (30–33). We previously found that integration into the genome requires a 22-bp attachment site (*att*) in LME-1 that is identical to a sequence on the *L. pneumophila* genome (33). Even though LME-1 is only integrated in a small proportion of sequenced *L. pneumophila* isolates, the chromosomal *att* site is present in all *L. pneumophila* isolates and *Legionella* species sequenced to date (Table S1), a level of conservation that suggests an important function. We report here that the LME-1 *att* site on the chromosome meets several criteria of the missing *dif* site in *Legionella*: size ∼30 bp, low GC-content, some degree of palindromicity, positioned close to the replication terminus, and in a non-coding region with varying flanking genes. We identify a single recombinase that is both necessary and sufficient for site-specific recombination at this site. Together, our results indicate that *L. pneumophila* uses a distinct single-recombinase system for CDR and that LME-1 is an IMEX of that system. We also report that *xerL* orthologs can be found across the Legionellales family. Lastly, we show that the *Coxiellaceae dif* site and *xerL* ortholog can functionally substitute for the *Legionella dif*/XerL system and propose that the migration of *C. burnetii* XerL off the chromosome has contributed to the stability of virulence plasmids in this pathogen.

## RESULTS

### The LME-1 attachment site is invariably close to the *Legionella* replication terminus

We previously determined that LME-1 integrates into the *L. pneumophila* genome via a 22-bp *att* site (33). The corresponding *att* site on the chromosome is contained within a broader 29-bp sequence that is conserved across all sequenced *L. pneumophila* strains and *Legionella* species (Fig. 1A, Table S1), suggesting that it performs an essential function. While the *att* site itself is conserved, its genomic neighborhood varies (33) (Table S1), indicating that the intrinsic function of this sequence may be unrelated to its flanking genes. In a search for attributes of the *att* site that may hint at its function, we analyzed its chromosomal location in all *L. pneumophila* strains with available genome sequences (Table S1). We found that despite the disparate gene neighborhoods, the location of the *att* site was roughly opposite the origin of replication in all cases. In fact, GC-skew analysis of each genome revealed that the *att* site was invariably close to the cumulative GC-skew maximum, which occurs at the site of replication termination (34) (Fig. 1B, Table S1). The *att* site is present within a small ∼8-kb window between 35 kb and 43 kb from the GC-skew inflection point (Table S1). This specificity of positioning extended to the *att* sites of other *Legionella* species, which are located between 89 bp and 79 kb from the point of GC-skew inflection (Fig. 1B, Table S1).

**Figure 1.**
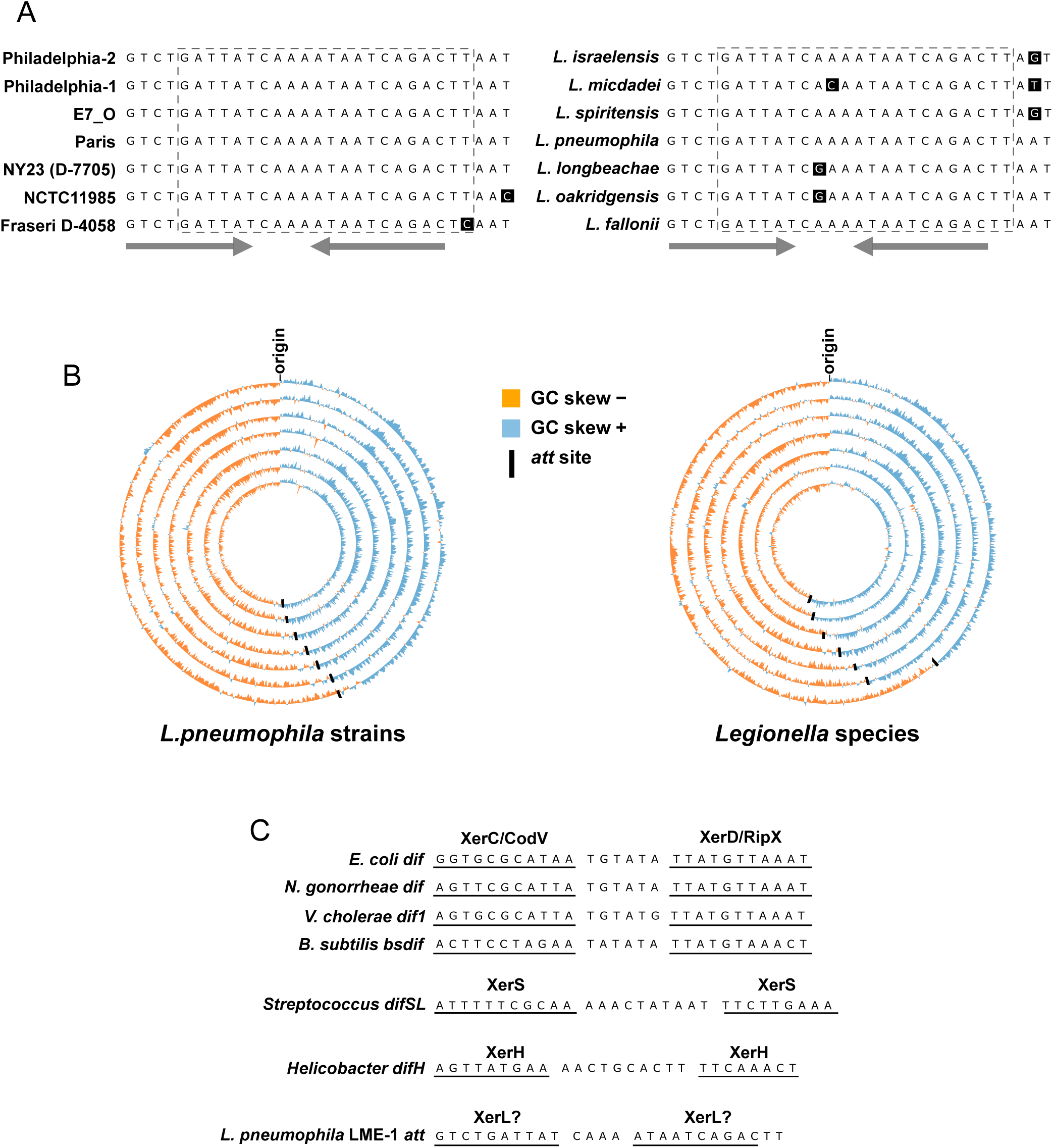
The DNA sequence hijacked by LME-1 is invariably proximal to the site of GC-skew inflection and resembles a *dif* site. **A)** Sequence alignment of the LME-1 *att* site region in different *L. pneumophila* strains (left panel) and other *Legionella* species (right panel). The dashed-line box denotes the 22-nt sequence that corresponds to the *att* site present on LME-1. The gray arrows indicate the inverted repeat portion of the sequence. **B)** Circos plots showing GC-skew across the chromosome of several *L. pneumophila* strains (left panel) and other *Legionella* species (right panel). Each ring corresponds to one genome and the black lines indicate the position of the *att* site. *L. pneumophila* strains inside to outside: Philadelphia-2, Philadelphia-1, E7_O, Paris, NY23 (D-7705), NCTC11985, Fraseri D-4058. *Legionella* species inside to outside: *L. israelensis*, *L. micdadei, L. spiritensis*, *L. pneumophila*, *L. longbeachae*, *L. oakridgensis*, *L. fallonii*. **C)** Alignment of the LME-1 *att* site with *dif* sites from bacterial species with established *dif*/Xer systems. The binding sites for XerC and XerD homologs (named CodV and RipX, respectively in *B. subtilis*) (10, 11, 18), XerS (64), and XerH (17, 65) are underlined.

The proximity of the LME-1 *att* site to the terminus region is reminiscent of bacterial *dif* sites, so we next compared the *att* site to the *dif* sites of several bacteria with established *dif*/Xer systems. In addition to its proximity to the terminus, the LME-1 *att* site shows several similarities to known *dif* sites, including (i) intergenic location (Table S1), (ii) size of ∼30 bp, (iii) low G+C content, and (iv) a short central region flanked by left and right arms with some degree of dyad symmetry (Fig. 1C). However, the LME-1 *att* site is distinguished from known *dif* sites by the perfect dyad symmetry of its 10-bp arms and the short 4-bp central region separating them. The arms do not show homology to any known Xer-binding motifs (Fig. 1C) (29).

### The *L. pneumophila dif* site supports highly efficient RecA-independent DNA recombination

The similarities between the chromosomal LME-1 *att* site (hereafter referred to as the *Legionella dif* site) and established *dif* sites suggest that it likely contains two binding sites for one or more tyrosine-type recombinases that catalyze site-specific recombination (3). To assess the ability of the *L. pneumophila dif* site to undergo site-specific recombination, we used an intermolecular recombination assay (Fig. 2A). In this assay, a non-replicative plasmid (pJB4648) containing a gentamicin-resistance marker and a 22-bp portion of the *dif* site (corresponding to the LME-1 *att* site) is transformed into *L. pneumophila* strain Lp02. If site-specific recombination occurs between the *att* site on the plasmid and the chromosomal *dif* site, the entire plasmid will be integrated into the genome and confer gentamicin resistance to the cell. We compared the number of transformants resulting from the *att*-containing plasmid to one with a control 22-bp sequence derived from the *L. pneumophila* genome near the *dif* site. To evaluate the efficiency of recombination supported by the *dif* site, we also included a plasmid containing 2200 bp of unrelated intergenic sequence from *L. pneumophila*, which can integrate into the genome via a single-crossover homologous recombination event. Considering that chromosomal *dif* site-specific recombination is known to be RecA-independent (13, 16, 35), we also assessed the integration of each of these plasmids in a *recA* deletion strain (Fig. 2B). To control for any strain-to-strain differences in overall transformation efficiency, we normalized the number of transformants for each plasmid to the number resulting from transformation with a replicative plasmid (pJB1806_GmR). The *att* site-containing plasmid integrated into the genome at a high efficiency that was 50-fold higher than that of the 2200-bp control. In contrast, the 22-bp control sequence did not generate any gentamicin-resistant recombinants, indicating that the number of integration events was lower than the detection limit of 1 CFU per ∼2 × 10^9^ cells. In the Δ*recA* strain, no transformants were detected for the 2200-bp control which relies on homologous recombination for integration, while the *att* site-containing plasmid integrated at a level similar to that in the wildtype strain (Fig. 2B). In both the wildtype and Δ*recA* strains, integration of the *att*-containing plasmid at the chromosomal *dif* site was confirmed by PCR-amplification of the *dif* region and Sanger sequencing (data not shown). These data indicate that, similar to other *dif* sites, site-specific recombination at the *L. pneumophila dif* site is efficient and RecA-independent.

**Figure 2.**
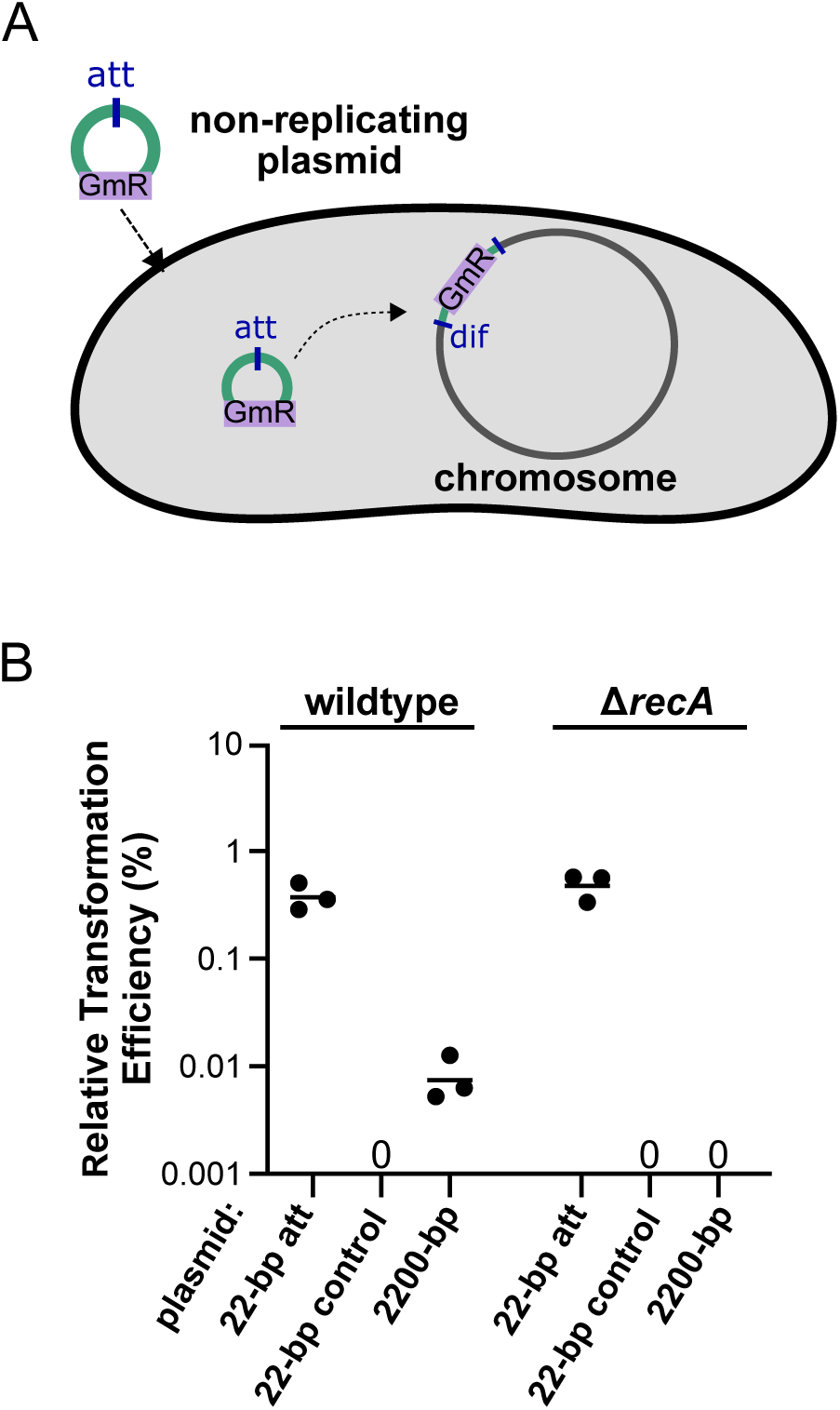
Site-specific recombination at the *dif* site is highly efficient and does not require RecA. **A)** Schematic representation of the intermolecular recombination assay used to quantify *att* site-specific recombination in *L. pneumophila*. A 5.5-kb non-replicating plasmid modified to include the 22-bp LME-1 *att* site (or control sequences) is transformed into *L. pneumophila* strain Lp02. If recombination between the plasmid *att* site and the chromosomal *dif* site takes place, the whole plasmid is integrated into the genome and gentamicin resistance is conferred to the cell. Gentamicin-resistant colonies are counted to quantify recombination. **B)** Transformation efficiency (relative to that of a replicative plasmid control) of a non-replicative plasmid containing the 22-bp *att* site, a 22-bp control sequence, or a 2200-bp intergenic region of the Lp02 genome, transformed into wildtype or Δ*recA* Lp02 strains. Each dot shows the value obtained from a single experiment, and each horizontal line represents the geometric mean of 3 independent experiments.

### *Legionella* Xer/*dif* is a single-recombinase system

We next aimed to identify the recombinase(s) that catalyzes recombination at the *Legionella dif* site. No orthologs of XerC, XerD, XerS, or XerH have been identified in *Legionella* using sequence homology (29). However, HHPred analysis (36) revealed that the proteins encoded by six *L. pneumophila* genes present in strain Lp02 (*lpg0980*, *lpg0981*, *lpg1070*, *lpg1085*, *lpg1867*, and *lpg2057*) show structural similarity to tyrosine-type recombinases (probability >99%). We therefore repeated our intermolecular recombination assay in strains containing individual deletions of these six genes, along with a strain lacking the *dif* site. As expected, deletion of the *dif* site resulted in no detectable integration of the *att* site-containing plasmid. Deletion of *lpg1867* also specifically abolished integration, while deletion of each of the other five recombinases had no significant effect (Fig. 3A). We also performed rescue experiments using expression plasmids to confirm that the loss of recombination at the *dif* site in the Δ*lpg1867* strain was due to loss of Lpg1867 protein function and not any secondary mutations acquired during strain generation. Expression of Lpg1867 in the Δ*lpg1867* strain resulted in near-wildtype levels of recombination (Fig. 3B). In contrast, no recombination was detected with an empty vector control or when a point mutant of Lpg1867 (Y387F) was expressed. The Y387F mutant is predicted to be catalytically inactive based on the equivalent catalytic tyrosine mutation in other tyrosine recombinases (10, 37, 38).

**Figure 3.**
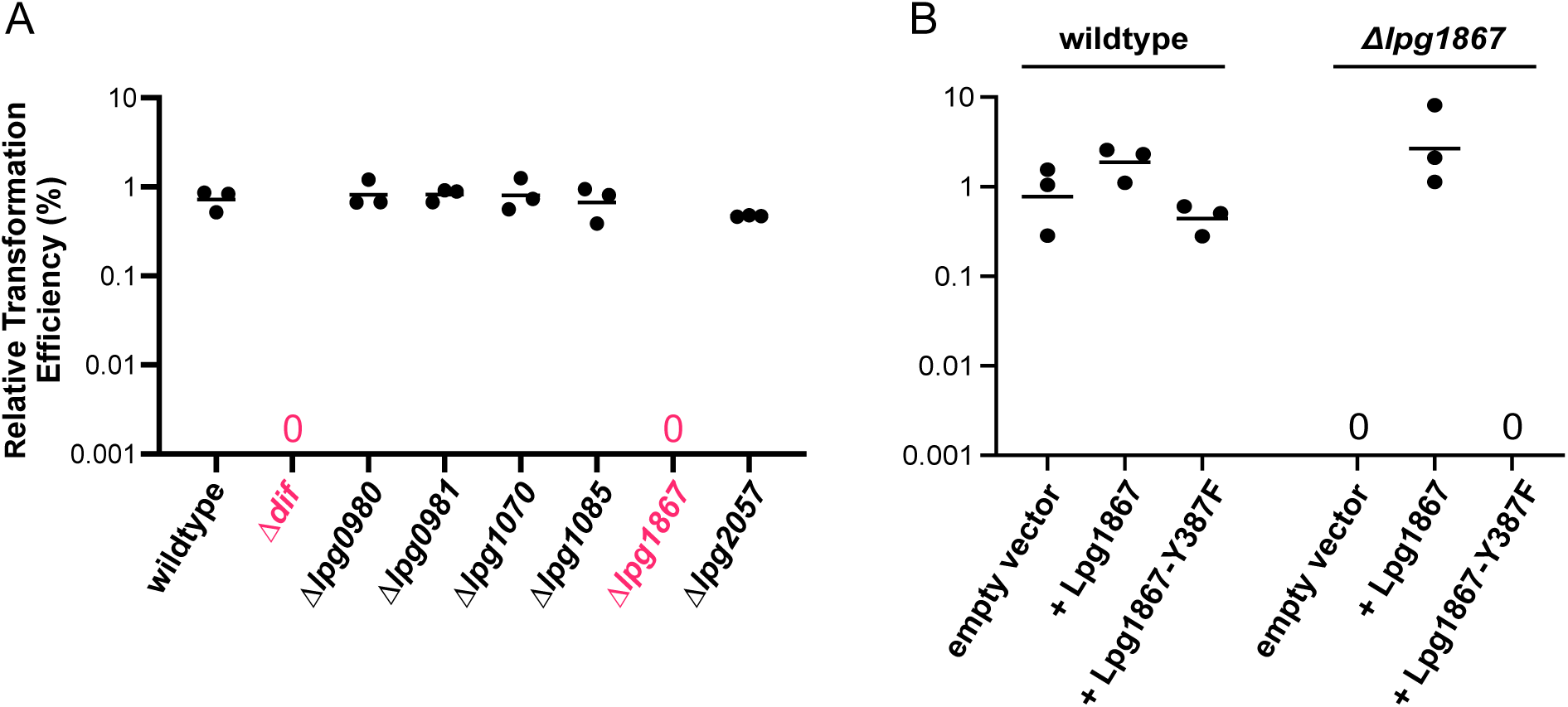
The *L. pneumophila* Xer is encoded by *lpg1867*. Dot plots showing levels of intermolecular recombination between the chromosomal *dif* site and a plasmid containing the 22-bp *att* site, as described for Figure 2. **A)** Intermolecular recombination in wildtype Lp02 and strains containing deletions of the *dif* site or one of six tyrosine-type recombinases. Shown in pink are the *dif*-deletion strain and *lpg1867*-deletion strain, which both displayed levels of recombination below the limit of detection. **B)** Intermolecular recombination in the wildtype and Δ*lpg1867* strains after transformation with plasmids expressing wildtype Lpg1867 or a predicted catalytic point mutant, Y387F.

We next asked whether one of the other five putative tyrosine recombinases might be involved along with Lpg1867 in recombination at the *dif* site, despite having no effect upon individual deletion. To address this, we generated two multiple-deletion strains: one with five putative recombinases deleted (all except *lpg1867*), and a pan-deletion strain with all six deleted. In our intermolecular recombination assay, integration of the *att*-containing plasmid was similar to wildtype levels in the 5-deletion strain and nonexistent in the pan-deletion strain (Fig. 4A). Overexpression of Lpg1867, but not Lpg1867-Y387F, again rescued the recombination defect (Fig. 4B).

**Figure 4.**
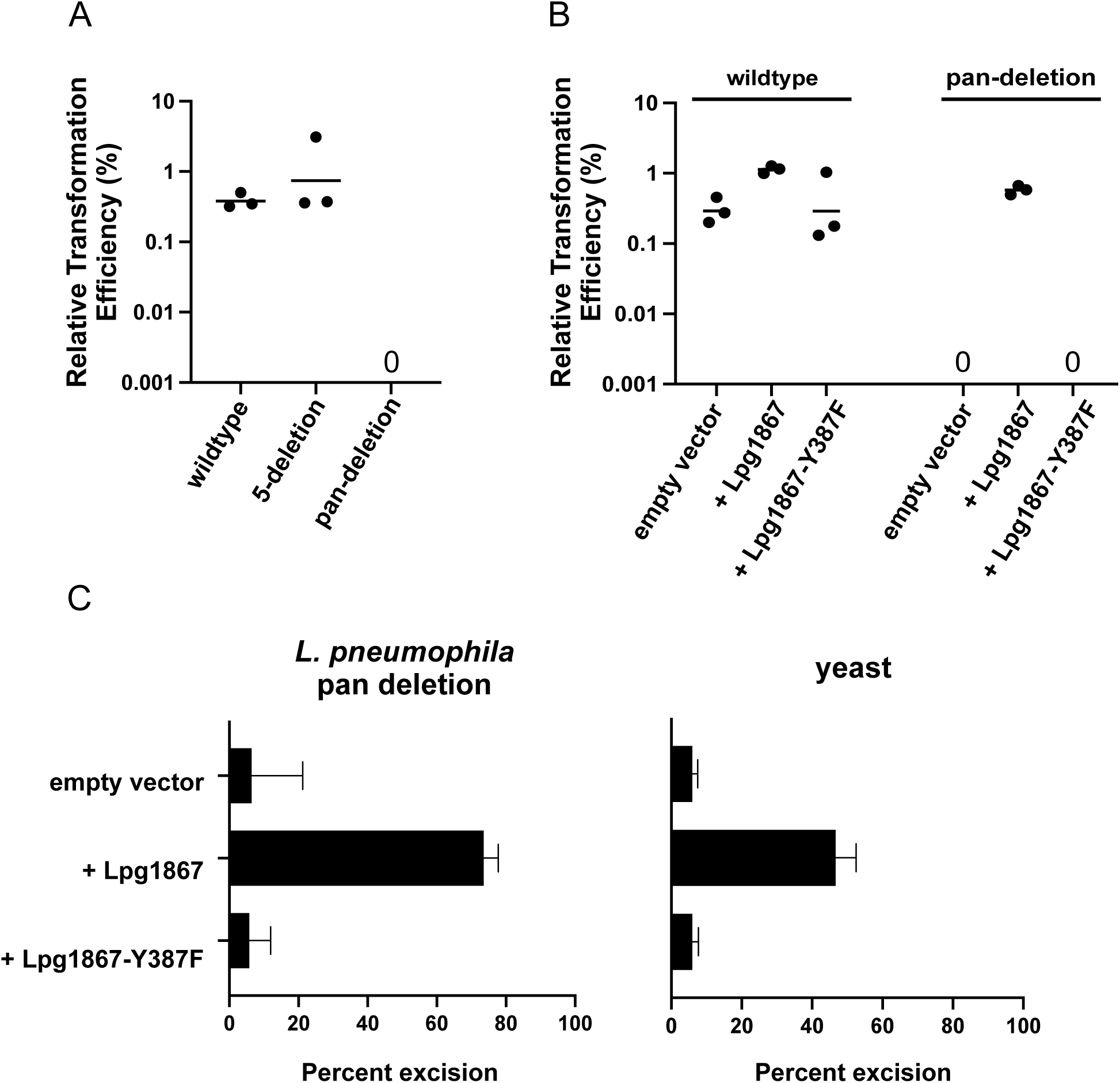
The *Legionella* CDR system involves a single recombinase. **A)** Intermolecular recombination in the wildtype Lp02 strain was compared to that in a 5-deletion strain (*lpg0980*, *lpg0981*, *lpg1070*, *lpg1085*, and *lpg2057* deleted) and a pan-deletion strain, with all six recombinases deleted (i.e, 5-deletion plus *lpg1867* deletion). **B)** Intermolecular recombination of the wildtype strain and pan-deletion strain containing the indicated expression plasmids. **C)** Left panel: Excision of a *dif* site-flanked kanamycin reporter in the Lp02 pan deletion strain to assess intramolecular recombination between *dif* sites. The proportion of kanamycin-resistant cells in the population after transformation with Lpg1867, the catalytic mutant Y387F, or empty vector control was quantified by plating on selective and nonselective media. Right panel: Excision of a *dif* site-flanked KanMX reporter in *S. cerevisiae* BY4741. Bars indicate the mean of three independent experiments, and error bars represent SEM.

As chromosome dimer resolution involves recombination between two *dif* sites on the same molecule, we also wanted to test whether Lpg1867 was sufficient to catalyze intramolecular recombination between two *dif* sites. To this end, we developed an intramolecular recombination assay by inserting a *dif* site-flanked kanamycin-resistance cassette at the normal *dif* locus of the Lp02 pan-deletion strain, such that recombination between the two flanking *dif* sites would result in loss of kanamycin resistance. To quantify excision of the reporter, we determined the proportion of kanamycin-resistant cells in the population after transforming the cells with wildtype or mutant Lpg1867. Overexpression of wildtype Lpg1867 in this strain resulted in a high level of cassette excision, while overexpression of the catalytic mutant resulted in an excision level similar to background level (Fig. 4C). Together, these data suggested that Lpg1867 does not require a partner recombinase for recombination at the *dif* site. However, to rule out the possibility that another unidentified recombinase or cofactor in *L. pneumophila* is also required, we performed a similar intramolecular excision assay in yeast. In this case, a *dif* site-flanked KanMX cassette was inserted at the HO locus of the *S. cerevisiae* BY4741 strain, and recombination between the two *dif* sites was assessed by quantifying loss of geneticin resistance (Fig. 4C). Overexpression of Lpg1867, but not Lpg1867-Y387F, again led to high levels of excision. Collectively, these data indicate that Lpg1867 is both necessary and sufficient to catalyze site-specific recombination at the *L. pneumophila dif* site.

### Deletion of *dif* or *lpg1867* induces filamentation and inhibits extracellular and intracellular growth

Loss of a functional chromosome dimer resolution system causes defects in cell division, which can result in a variety of phenotypes including slow growth and filamentation (the *dif* site was named for its <u>d</u>eletion-induced filamentation phenotype). Consistent with reported findings for *dif* or *xer* deletions in other species (11, 13, 14, 18, 39, 40), we observed that loss of *dif* or Lpg1867 in *L. pneumophila* led to filamentation in a subpopulation of cells (Fig. 5A). Notably, and consistent with a reduction in chromosome dimer formation in the absence of RecA (2, 41), we observed a decrease in filament formation upon deletion of *recA* in the Δ*lpg1867* background (Fig. 5A).

**Figure 5.**
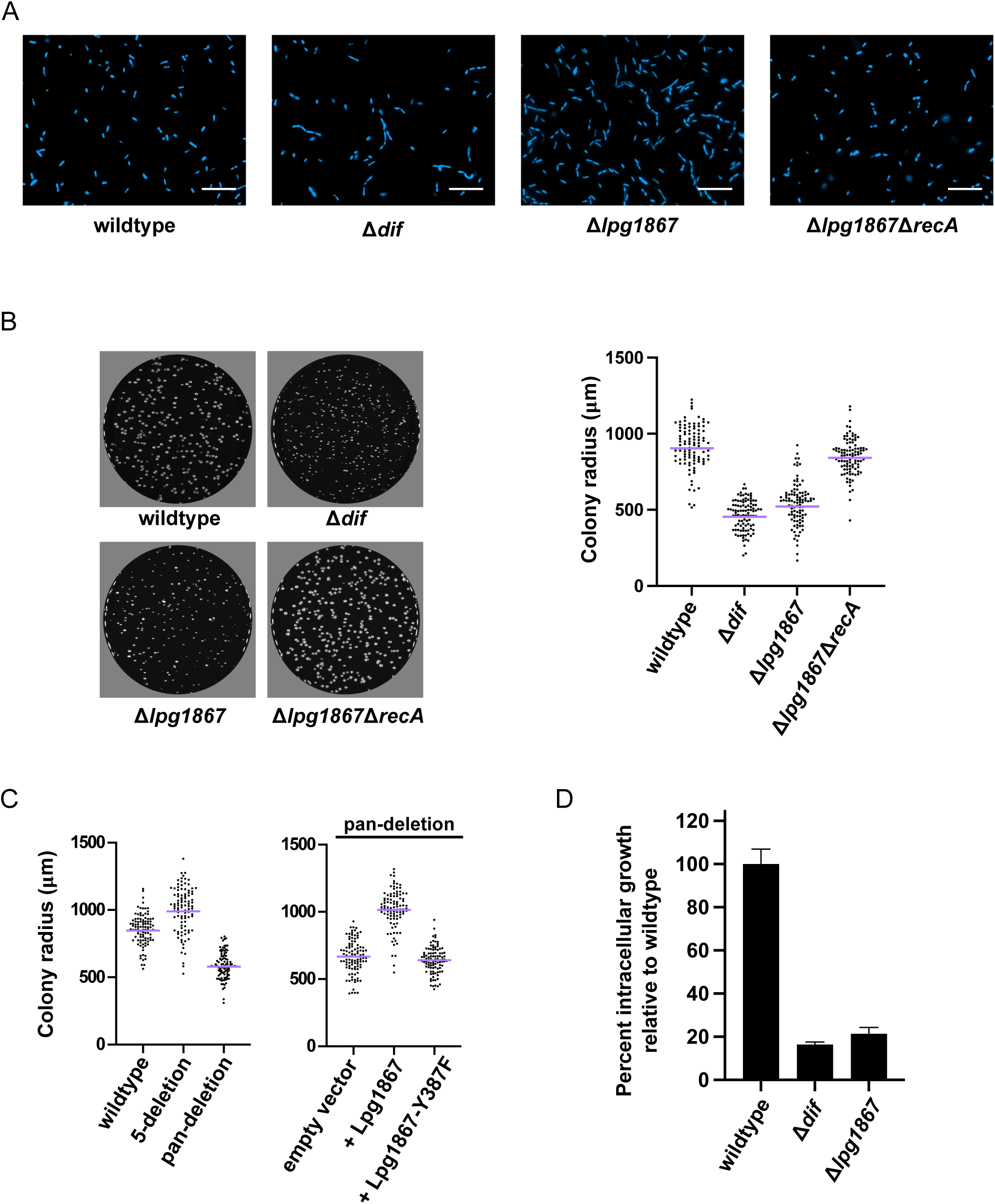
*dif- and lpg1867-*deletion strains exhibit filamentation and altered growth on solid media and during infection of host cells. **A)** DAPI-stained cells of the indicated *L. pneumophila* strains, showing filamentation upon deletion of the *dif* site or *lpg1867* and rescue when *recA* is deleted in the Δ*lpg1867* background. The scale bars represent 10 μm. **B)** Left panel: CYET spread plates after 4 days of growth at 37℃ with visible differences in colony size. Right panel: Each colony radius is plotted as a separate data point, with the geometric mean shown by the purple line. **C)** Distribution of colony sizes in the 5-deletion strain and pan-deletion strain (left panel) and in the pan-deletion strain expressing wildtype or mutant Lpg1867 (right panel). Data are plotted as in B. **D)** Intracellular growth of wildtype and deletion strains in U937 macrophages. The data are plotted as the fold increase in colony-forming units recovered after lysis from host cells at 2 h and 48 h post infection, relative to wildtype Lp02. Data show the average of three replicates for each strain from a representative experiment. Error bars represent SEM.

Deletion of *dif* or *lpg1867* also results in a visible reduction in colony growth on solid media (Fig. 5B). Again, this defect is alleviated upon additional deletion of *recA* in the *Δlpg1867* background (Fig. 5B), consistent with the growth inhibition being caused by an inability to resolve chromosome dimers. The slow-growth phenotype is also present in the recombinase pan-deletion strain, but not the 5-deletion strain, and the defect is rescued by expression of wildtype Lpg1867 but not the catalytic mutant (Fig. 5C), mirroring the effects of these deletions on recombination (see Fig. 4A,B). Importantly, the *dif* and *lpg1867* deletions also result in diminished growth in human U937 macrophages (Fig. 5D), indicating that growth within host cells is also impacted by disruptions to *dif*/Xer activity. Taken together, these data support a model in which a single site-specific recombinase, Lpg1867 (which we designate here as XerL), catalyzes recombination between two *dif* sites to resolve chromosome dimers in *Legionella*.

### LME-1 stability is modulated by the formation of a modified downstream *dif* site

We have previously shown that LME-1 integrates into the *L. pneumophila* chromosome at what we now know to be the *dif* site (33). LME-1 is stably integrated into the genome of str. Murcia-4983 (only excised in ∼1% of the population), despite being flanked by two *att* sites (33). The sequence of the LME-1 *att* site is identical to the *L. pneumophila dif* site, but only encompasses 22-bp of it. Recombination between this 22-bp sequence and the chromosomal *dif* site results in *att*-site duplication, which generates an intact *dif* site upstream and a modified *dif* site downstream of LME-1 that contains three substitutions (Fig. 6A). We next asked whether these modifications to the downstream *dif* site contribute to the stability of the integrated form of LME-1. We modified our excision assay to use the kanamycin-resistance cassette as a proxy for LME-1, and determined the effect of adding one or more of the LME-1 *att* substitutions to the downstream *dif* site (Fig. 6B). We found that adding all three substitutions to the downstream *dif* site (to mimic integrated LME-1) resulted in markedly reduced excision compared to the reporter flanked by two wildtype *dif* sites, while one or two substitutions resulted in an intermediate level of excision (Fig. 6B). These data suggest that the short 22-bp *att* site of LME-1 likely contributes to the stability of its integrated form by preventing it from being flanked by two intact *dif* sites.

**Figure 6.**
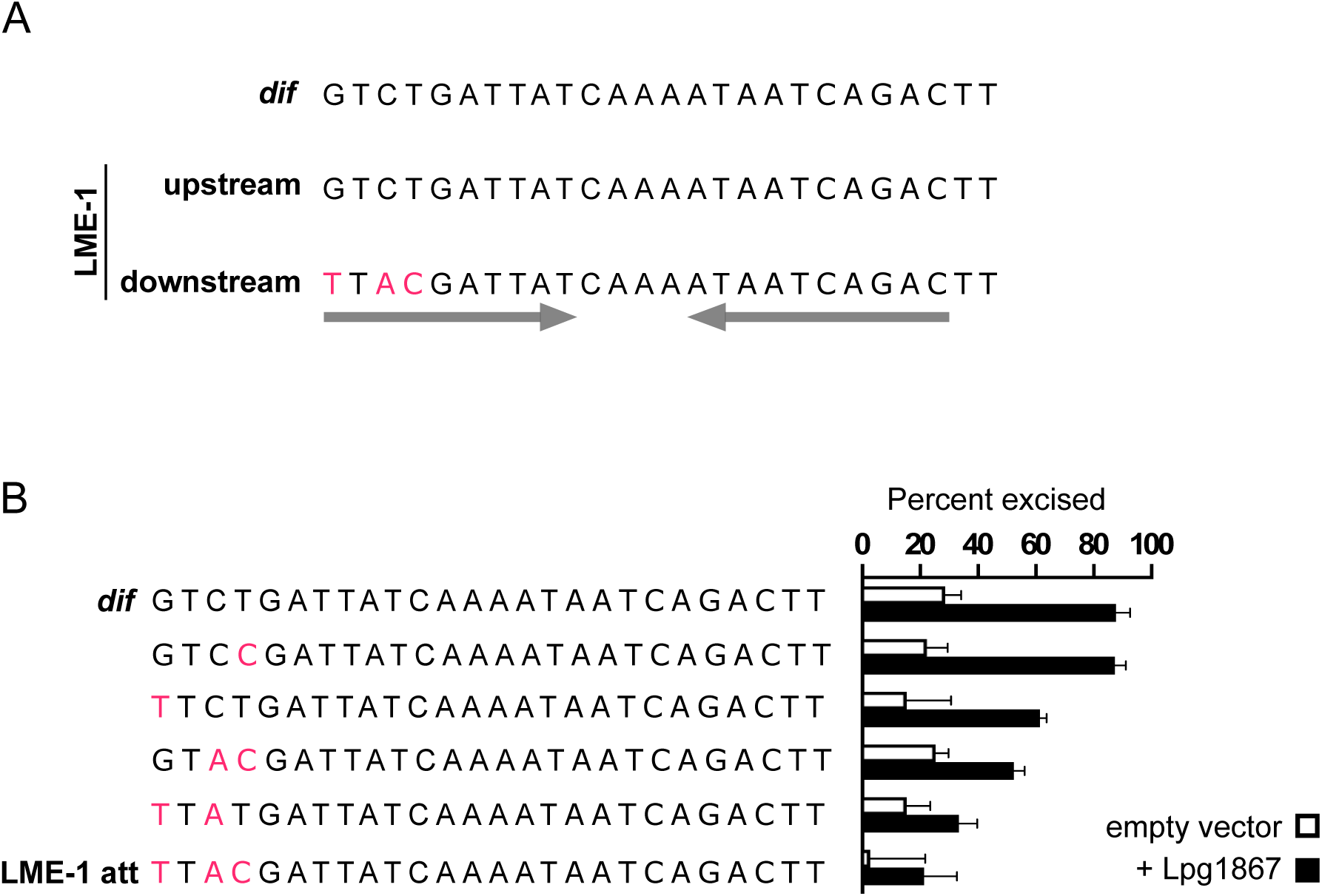
The sequence of the LME-1 *att* site contributes to its low level of excision, which increases upon Lpg1867 (XerL) overexpression. **A)** Comparison of the wildtype *dif* site in strain Murcia-4983 and the sequences flanking LME-1 resulting from *att* site duplication upon integration into the chromosome. The upstream copy of the *dif* site remains intact, but the downstream copy has three nucleotide differences (pink letters). These differences are within the inverted repeat regions of the *dif* site (denoted by the gray arrows), which presumably contain binding sites for Lpg1867 (XerL). B) A kanamycin-resistance reporter cassette flanked by a wildtype *dif* site upstream and the indicated sequence downstream was introduced into the Lp02 Δ*lpg1867* strain. Excision of the reporter (indicating recombination between the two flanking sites) was quantified by plating on selective and nonselective media after transformation with empty vector or a plasmid expressing Lpg1867.

### Widespread distribution of *xerL* orthologs in *Legionella* and *Coxiella*

Our discovery of the *Legionella dif*/Xer pathway, which had been missed in homology-based searches due to divergence from known CDR components, led us to next examine other bacteria in which no *dif*/Xer system has been identified to date. One possibility is that several species have XerL-like systems that were missed for the same reasons that they were missed in *Legionella*. To explore this possibility, we performed BLASTp and tBLASTn analyses using the *L. pneumophila* XerL protein as a query. These analyses identified several additional XerL orthologs across the order Legionellales, which includes *Legionella* sp. as well as the select agent, *Coxiella burnetii* (Fig. 7).

**Figure 7.**
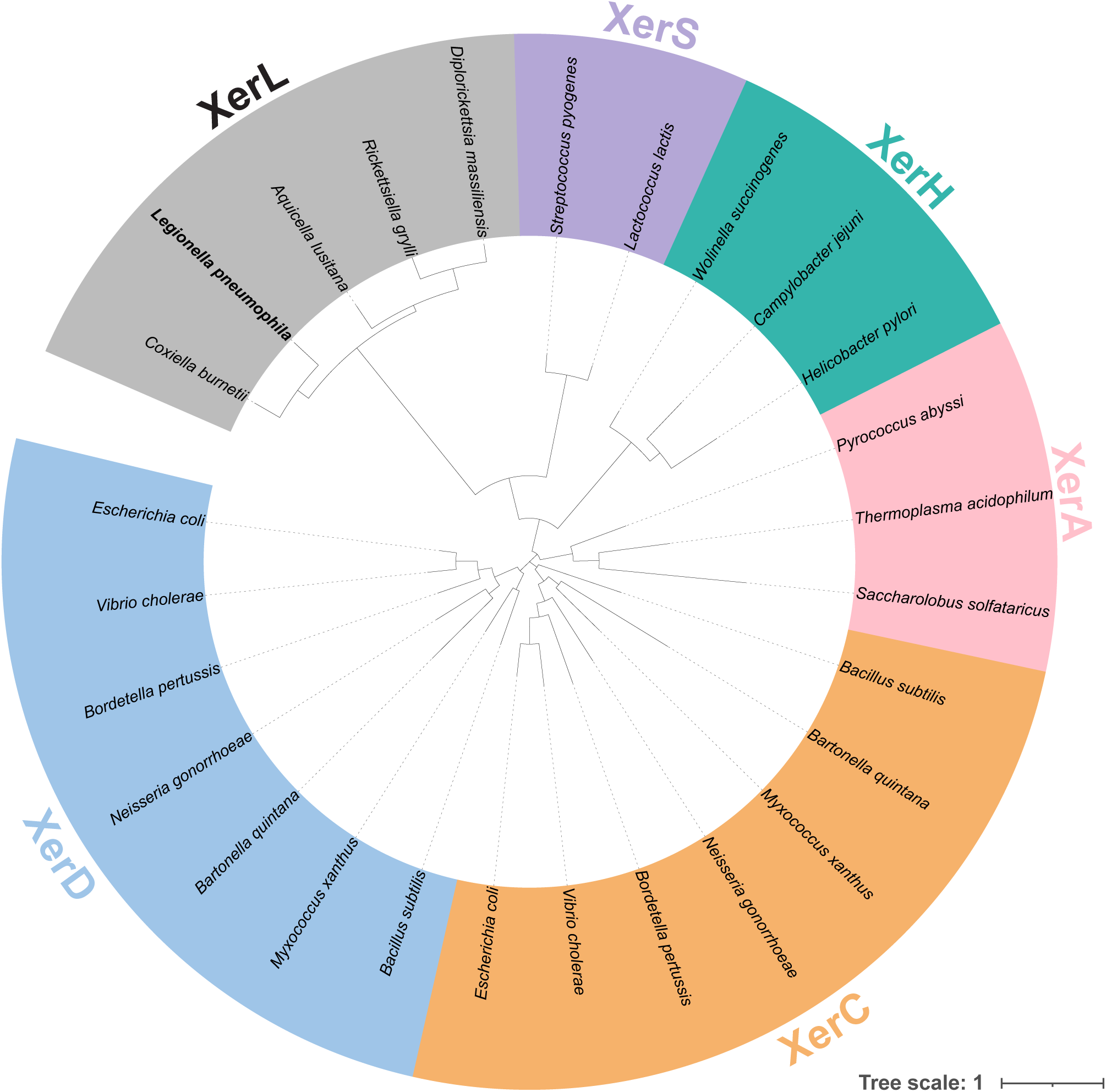
Phylogenetic analyses suggest that XerL forms a distinct clade from other Xer recombinases. XerL from *L. pneumophila* was used as a query to search for putative XerL orthologs using BLASTp and tBLASTn. The amino acid sequences of recombinases from different Xer groups were aligned with MUSCLE, with one representative per genera included for the XerL-like recombinases. An unrooted phylogeny was generated from the alignment using FastTree. Xer group designation is denoted with colored labels as indicated. The scale bar indicates the number of amino acid substitutions per site.

Phylogenetic analysis of the XerL-like recombinases indicates widespread distribution across Legionellales (Fig. S1). The amount of sequence divergence between XerL and the rest of the Xer protein family likely explains how this machinery has been missed by prior analyses.

### The *Coxiella burnetii xerL* ortholog is encoded on a virulence plasmid

Despite the apparent vertical inheritance of *xerL* given its restriction within the order Legionellales, the *Coxiella burnetii* orthologs of *xerL* (hereafter *xerL*_Cb_) reside not on the circular chromosome, but on a large ubiquitous plasmid that is critical for virulence (42–45). While Xer/*dif* systems have established roles in resolving aberrant plasmid dimers, these are the only Xer sequences within the *C. burnetii* genome. This raises the question as to whether this plasmid-based XerL functions to resolve plasmid dimers, chromosome dimers, or both.

To search for putative *dif* sites within the *Coxiella* genome, we performed a sequence similarity (BLASTn) search using the 29-nt *L. pneumophila dif* site. We identified a short sequence of 30 nucleotides on the virulence plasmid immediately adjacent to XerL_Cb_ (Fig. 8A). Using this second sequence as a query, we identified an additional site on the chromosome (Fig. 8A). This chromosomal sequence diverges from the plasmid site yet has several features consistent with a functioning *dif* site. Like the *L. pneumophila dif* sequence, it is palindromic, highly conserved across isolates, and has no apparent coding potential.

**Figure 8.**
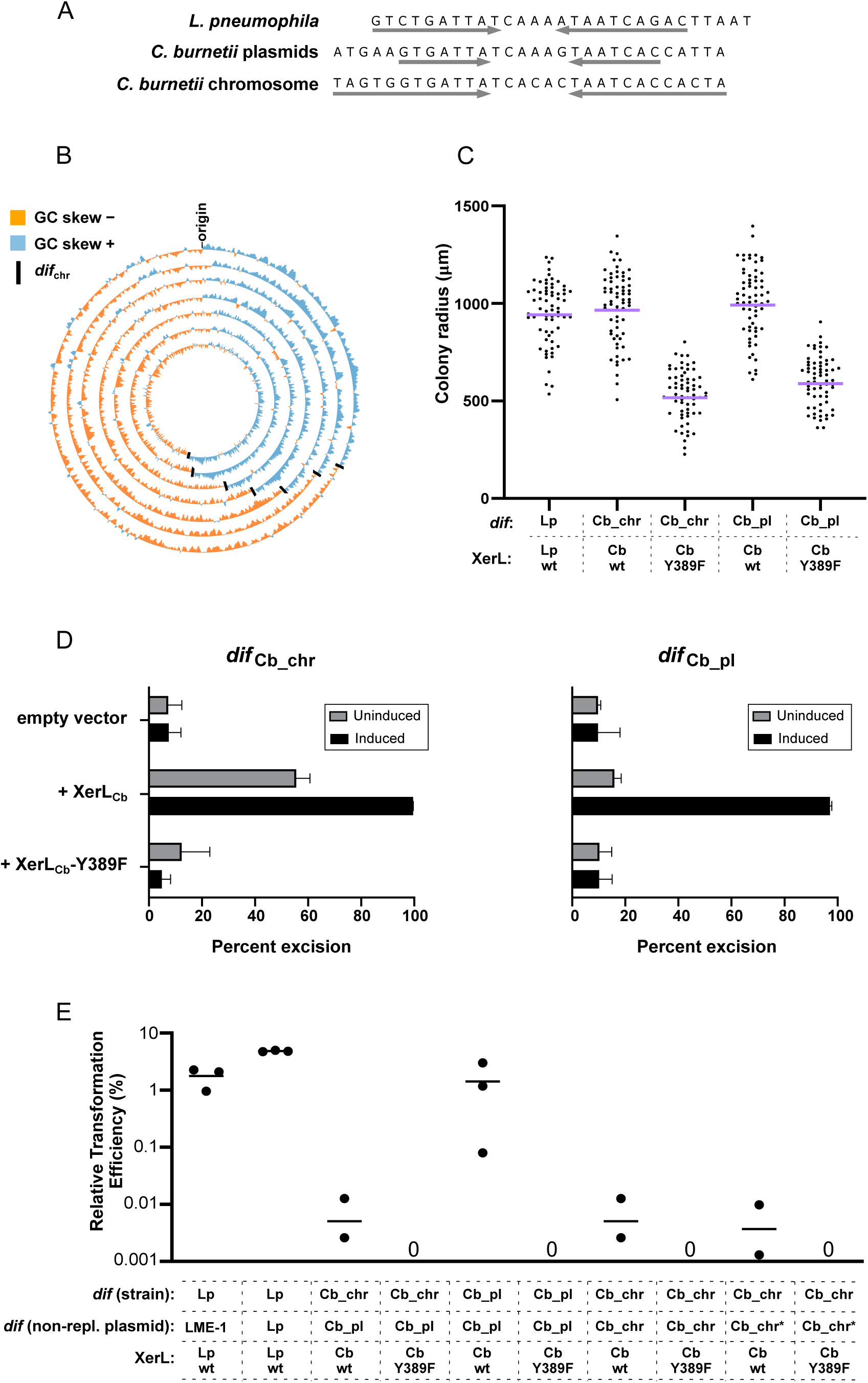
The *C. burnetii dif* site and XerL can functionally replace endogenous *dif*/Xer in *L. pneumophila*. **A)** Sequence alignment of the *L. pneumophila dif* site with the plasmid and chromosomal *dif* sites of *Coxiella burnetii*. The gray arrows indicate the inverted repeat portion of the sequence. **B)** Circos plot showing GC-skew across the chromosome of several *C. burnetii* strains. Each ring corresponds to one genome and the black lines indicate the position of the *dif* site. *C. burnetii* strains, inside to outside: 3262, Heizburg, Dugway FJ108-111, RSA 493, CbuG_Q212, Schperling, CbuK_Q154. **C)** Strains were generated in which the endogenous *L. pneumophila dif* was replaced by the *C. burnetii* chromosomal *dif* site (*dif*_Cb_chr_) or plasmid *dif* site (*dif*_Cb_pl_), and the endogenous XerL was replaced by *C. burnetii* XerL or a predicted catalytic point mutant (Y389F). Colony sizes for each strain were quantified after 4 days of growth at 37℃ on CYET spread plates. Each colony radius is plotted as a separate data point, with the geometric mean shown by the purple line. **D)** Intramolecular recombination assay performed as described for Figure 4C (uninduced) and with an additional overnight incubation with 100 ng/mL anhydrotetracycline (induced). Percent excision of a kanamycin reporter cassette flanked by the indicated *C. burnetii* chromosomal *dif* site (*dif*_Cb_chr_; left panel) or plasmid *dif* site (*dif*_Cb_pl_; right panel). **E)** Intermolecular recombination assay performed as described for Figure 2, showing the relative transformation efficiency of the indicated non-replicative plasmids in strains containing the indicated *dif* site and *xerL*. Each dot shows the value obtained from a single experiment, and each horizontal line represents the mean of 3 independent experiments. The Cb_chr* *dif* site is a truncated version of the chromosomal dif site, with truncations made to mimic the shorter 22-bp LME-1 *att* site version of *dif*_Lp_.

To further investigate the potential role of this chromosomal site as a functional *dif* site, we examined its location relative to the point of inflection of GC skew (replication terminus). As we observed for *dif*_Lp_, the *Coxiella burnetii* chromosomal site is always located opposite the predicted origin of replication and close to the cumulative GC-skew maximum (Fig. 8B, Table S2). Taken together, this sequence not only shares homology to *dif*_­_, but maintains several hallmarks of a *dif* site involved in chromosome dimer resolution. Hereafter, we will refer to it as *dif*_Cb_.

### The *Coxiella* XerL/*dif* system functionally complements a Δ*xerL*/*dif* mutant in *Legionella*

Having identified putative *dif* sites and *xerL* orthologs in *Coxiella burnetii*, we next wanted to assay their potential for chromosome dimer resolution. Considering that colony size correlates with *dif*/Xer function in *L. pneumophila* (Fig. 5), we generated a strain in which the endogenous *dif* site and *xerL* gene were replaced by the *C. burnetii dif* site and *xerL*, and assessed growth on solid media. If the *C. burnetii dif*/XerL were not able to resolve dimers in their surrogate chromosome, then we would expect small colony sizes similar to the Δ*dif* or Δ*xerL* strains. However, the colony size distribution of the strain containing XerL_Cb_ and the *C. burnetii* chromosomal *dif* site (*dif*_Cb_chr_) was almost identical to that of wildtype *L. pneumophila* (Fig. 8C), while a strain encoding the predicted catalytic mutant XerL-Y389F displayed the small colony sizes associated with nonfunctional *dif*/Xer (Fig. 8C). This suggests that despite being encoded on the plasmid, XerL_Cb_ may catalyze recombination at *dif*_Cb_chr_ to resolve dimers of the *Coxiella* chromosome. Interestingly, despite containing several substitutions (Fig. 8A) the *Coxiella* plasmid *dif* site (*dif*_Cb_pl_), in concert with XerL_Cb_, is also able to complement the small colony phenotype (Fig. 8C). Similarly, in our intramolecular recombination assay, the reporter cassette was excised when flanked by either *dif*_Cb_chr_ or *dif*_Cb_pl,_ although excision for the cassette flanked by *dif*_Cb_pl_ required overnight induction of XerL_Cb_ (Fig 8D).

Together, these findings suggest that XerL_Cb_ may resolve both plasmid and chromosome dimers in *Coxiella*, with the resolution of chromosome dimers uniquely dependent on plasmid-encoded XerL. XerL is encoded by all four virulence plasmids described in *C. burnetii* and no additional Xer orthologs can be found on the chromosome. This suggests that chromosome dimer resolution and virulence plasmid maintenance are genetically linked in *C. burnetii*, with plasmid loss leading to loss of the chromosome dimer resolution pathway. Consistent with this model, we note that previously described instances of plasmidless strains – in which a subset of plasmid-like sequences are instead integrated onto the chromosome (46–48) – contain *xerL* within the integrated sequence. Our findings suggest that such integrants are likely selected for based on their ability to escape the fitness costs that a plasmidless strain would otherwise incur. Despite the sequence similarity between the *dif* sites located on the plasmid and chromosome, and their shared dependence on XerL, the integration of plasmid-like sequences in the plasmidless strains is not at the chromosomal *dif* site. In fact, the plasmid fragments are located ∼1 Mb from the terminus region, indicating that acquisition of the plasmid-like sequences did not originate with an IMEX-like plasmid integration event. To see if this phenomenon would recapitulate in our intermolecular recombination assay, we measured integration of a non-replicative plasmid containing *dif*_Cb_pl_ into the genome of a strain containing *dif*_Cb_chr_ and found it to be almost 300-fold lower than when the genome contained *dif*_Cb_pl_ (Fig. 8E). Surprisingly, the efficiency of integration into the *dif*_Cb_chr_-containing genome was also very low when the non-replicative plasmid contained the identical *dif*_Cb_chr_ (Fig. 8E). One possible explanation is that highly efficient recombination between the two *dif*_Cb_chr_ sites leads to rapid re-excision after integration, resulting in no visible gentamicin-resistant colonies. However, when we used a version of *dif*_Cb_chr_ with a shortened region of dyad symmetry, which is known to increase stability of integration for the LME-1 *dif* mimic (see Fig. 6), transformation efficiency remained very low (Fig. 8E). These findings suggest that either shortened dyad symmetry does not stabilize *dif*_Cb_chr_ or the chromosomal *dif* site itself might be recalcitrant to invasion. The latter notion is consistent with the fact that no sequenced *Coxiella burnetii* genomes contain virulence plasmids as chromosomal integrants at the *dif* site.

## DISCUSSION

The discovery of the *L. pneumophila* CDR system has solved two mysteries of the pathogen’s biology: the function of the conserved DNA sequence that is hijacked by LME-1, and how *Legionella* species are able to resolve chromosome dimers despite having no homologs to known *dif* sites and Xer recombinases. Similarly, no *dif*/Xer components had been identified previously in members of the other Legionellales family, *Coxiellaceae* (29), which includes the human pathogen *Coxiella burnetii*. It is rare that you uncover an overlooked, essential pathway in a pathogen. We have identified one that is conserved across an entire order. Other extensively studied dimensions of *Legionella* biology (e.g. translocated effectors) are not strong candidates for drug targets due to their redundancy. In contrast, we show that disrupting one protein in this pathway (XerL) is sufficient to dramatically restrict the growth of the pathogen.

With no apparent *dif* or Xer homologs in *Legionella*, the CDR system was instead unearthed through our investigation of LME-1 and its attachment site. In turn, the discovery of the *Legionella dif* site and XerL has allowed us to explore new LME-1 biology. Our findings are consistent with LME-1 belonging to the IMEX class of integrative mobile elements, which exploit the host CDR machinery for integration and in some cases excision (5, 28). The mechanistic details of LME-1 integration and excision, and how they relate to the strategies of other IMEXs, are interesting topics of future study. One intriguing aspect of LME-1 is the stability of its integrated form. Our reporter excision assay showed that the modified *dif* site and wildtype *dif* site that flank the integrated LME-1 recombine at a very low level, and that the stability of integration is influenced by the three nucleotide substitutions present in the modified form. Another stably integrated IMEX, the gonococcal genomic island (GGI) of *Neisseria gonorrhoeae*, generates a similarly modified *dif* site (*dif*_GGI_) at one end of its integrated form, while the wildtype *dif* (*dif*_Ng_) remains intact at the other (23). The *Neisseria* CDR system uses XerC and XerD homologs, and recombination takes place when the FtsK DNA translocase pauses at the *dif/*Xer complex and activates XerD. Interestingly, FtsK does not stop at the *dif*_GGI_-XerC/D complex, but rather translocates through it, which likely leads to disassembly of the complex and precludes recombination between the *dif_NG_ and dif_GGI_* (23). It is possible that this same FtsK-dependent process contributes to the stability of the integrated form of LME-1, although we have yet to determine whether XerL requires FtsK for activation. Our future investigative priorities include determining the involvement of FtsK in the *L. pneumophila* CDR system and the stability of its associated IMEX, LME-1.

The *Legionella* and *Coxiella dif*/Xer systems are distinct in several ways from those characterized previously in other bacterial species. The *dif* site arms consist of long inverted repeats (10-bp and 12-bp, respectively) with no interruptions, whereas other *dif* sites show only partial dyad symmetry between the two arms, even for single-recombinase systems. Additionally, in *L. pneumophila*, *dif* and *xerL* are separated by >400 kb in the genome, whereas in the other bacterial single-recombinase systems (*i.e*. *dif*_SL_/XerS of *Streptococci* and *Lactococci* and *dif*_H_/XerH of *Campylobacter* and *Helicobacter*), the *dif* site is always near the *xer* gene, indicating they may have been acquired as a single module (3, 16, 17, 29). The location of the *C. burnetii xerL* ortholog on its virulence plasmid is also unusual. While many plasmids use Xer machinery for their own dimer resolution, this is the first instance we are aware of where a plasmid-based Xer recombinase is necessary for chromosome dimer resolution.

Given the general importance of chromosome dimer resolution to bacterial fitness and the extent to which intracellular growth is compromised in *L. pneumophila* Δ*xerL* or *dif* mutants, we anticipate that plasmid loss in *Coxiella* incurs a significant fitness cost through the loss of the essential recombinase. Consistent with this model, isolates that have lost their plasmid appear to maintain *dif*/XerL activity by including *xerL* within the fragments of plasmid sequence found to be integrated on the chromosome. One interpretation of these results is that integrants that do not maintain *xerL* chromosomally are rapidly selected against and not recovered. In light of our data, it will be important to distinguish which phenotypes previously associated with directed plasmid-loss (44) reflect virulence-specific defects (associated with loss of specific virulence factors, such as effectors) and which phenotypes result from the loss of *dif*/XerL machinery.

## MATERIALS AND METHODS

### Strains and plasmids

The *L. pneumophila* Lp02 strain was used as the background for all *L. pneumophila* strains generated in this study. The *L. pneumophila* Murcia-4983 strain, which harbors LME-1, is an environmental isolate collected during the 2001 outbreak in Murcia, Spain (49). All *L. pneumophila* strains were grown at 37℃ in N-(2-acetamido)-2-aminoethanesulfonic acid (ACES) buffered yeast extract supplemented with 100 ug/mL thymidine (AYET) and on charcoal AYET (CYET) agar plates.

Deletion and mutant strains were made using a scar-free suicide-cassette method described recently (50) with some minor modifications. Briefly, linear DNA containing a mazF-KanR cassette flanked by ∼1.5-kb homology arms was introduced into the parental strain via natural transformation. Cells in which the mazF-KanR cassette had been integrated via homologous recombination were selected by plating on media containing kanamycin. As mazF gene expression is under an IPTG-inducible promoter, colonies can be screened on kanamycin- and IPTG-containing plates to identify cells that are IPTG-sensitive and kanamycin-resistant. In a second natural transformation, linear DNA containing only the homology arms with the deletion/mutation was introduced, and cells in which the cassette was excised were selected on plates containing IPTG. Colonies were then screened to identify those that were IPTG-resistant and kanamycin-sensitive, and the deletion was verified by PCR screening and Sanger sequencing.

The reporter strains used for the intramolecular recombination (reporter excision) assays were generated by introducing linear DNA containing the *dif*-KanR-*dif* cassette (flanked by ∼1.5 kb homology arms around the chromosomal *dif* site) into the Lp02 Δ*lpg1867* strain or pan-deletion strain. Strains containing the cassette were selected on plates containing kanamycin, and the correct integration of the cassettes was verified by Sanger sequencing. Similarly, for the yeast-based excision assay, linear DNA containing the *dif*-KanMX-*dif* cassette (flanked by homology arms around the HO locus) was transformed into *S. cerevisiae strain* BY4741 using the high efficiency PEG/LiAc method (51). Integrants were selected on YPD agar plates containing geneticin and verified by Sanger sequencing.

The pJB4648-based plasmids used for the intermolecular recombination assays were constructed by cloning the 22-bp *att* site sequence, a control 22-bp sequence, or 2200 bp of intergenic Lp02 sequence into pJB4648 digested with XhoI and ApaI. The gentamicin-resistant plasmid pJB1806-GmR was generated by first amplifying the gentamicin resistance gene and promoter from pJB4648 and cloning it in place of the chloramphenicol resistance gene in plasmid pJB1806. Protein expression plasmids used for the recombination and colony size assays were made by cloning the *lpg186*7 gene (wildtype or modified to encode the Y387F mutation) into the multiple cloning site of pNT562 (52). The tet-on-pJB1806-GmR-based expression plasmids used for the reporter excision assays were generated by replacing the *lacIq* gene and tac promoter in pJB1806-GmR with the *tetR* gene and tetracycline-responsive promoter from pLJR965 (53) and then cloning in the wildtype or mutant Lpg1867 ORF immediately after the promoter, as described by Rock and colleagues (53).

### GC-skew analysis

The cumulative GC-skew maximum value for each *Legionella* genome (Table S1) or *C. burnetii* genome (Table S2) was determined using the GenSkew java app (https://genskew.csb.univie.ac.at) with default settings. The whole-genome GC-skew profiles and *dif* site locations for seven *L. pneumophila* isolates, seven *Legionella* species, and seven *C. burnetii* strains, selected to show a range of cGC-skew maxima locations relative to the origin, were plotted using Circos (54). The GC-skew profiles of these genomes were normalized for genome length. The selected *L. pneumophila* isolates and *Legionella* species are: *L. pneumophila*: Philadelphia_2 (GenBank accession number CP015929.1), Philadelphia_1 (AE017354.1), E7_O (CP015954.1), Paris (CR628336.1), NY23 D-7705 (CP021261.1), NCTC11985 (LT906452.1), and subsp. Fraseri strain D-4058 (CP021277); *Legionella* species: *L. israelensis* L18-01051 (CP041668.1), *L. micdadei* (LN614830.1), *L. spiritensis* NCTC11990 (LT906457.1), *L. sainthelensi* NCTC12450 (LR134178.1), *L. longbeachae* NSW150 (FN650140.1), *L. oakridgensis* NCTC11531 (LR134286.1), and *L. fallonii* LLAP-10 (LN614827.1). The selected *C. burnetii* isolates are: 3262 (CP013667.1), Heizburg (CP014561.1), Dugway FJ108-111 (CP000733.1), RSA 493 (AE016828.3), CbuG_Q212 (CP001019.1), Schperling (CP014563.1), and CbuK_Q154 (CP001020.1).

### Recombination assays

For the intermolecular recombination (plasmid integration) assays, each strain was transformed with a pJB4648-based plasmid containing the indicated *att*, *dif*, or control sequences as indicated. The strains were also transformed with the replicative plasmid pJB1806-GmR to control for strain-to-strain differences in general transformation efficiency. Plasmid concentrations were determined using a Quant-iT PicoGreen dsDNA kit (Invitrogen), and 200 ng of each plasmid was electroporated into 2 ODU of *L. pneumophila* cells as described previously (55). After an 8-h recovery in liquid AYET media shaking at 37℃, dilutions of each sample were plated onto CYET plates supplemented with 15 μg/mL gentamicin and incubated at 37℃ for 4 days. Colonies were counted using GeneTools analysis software (Syngene), and the results were plotted using GraphPad Prism version 9.2.0.

For the intramolecular recombination (excision) assay in *Legionella*, strains containing the kanamycin-resistance cassette were transformed with tet-on-pJB1806-GmR containing wildtype Lpg1867, Lpg1867-Y387F, XerL_Cb_, XerL_Cb_-Y389F, or empty vector by electroporation as described above. Transformants were selected on CYET plates containing both gentamicin and kanamycin. The transformant colonies were then collected in AYE and spread onto nonselective plates and plates containing kanamycin to quantify the proportion of the population that had lost kanamycin resistance. For the intramolecular recombination (excision) assay in yeast, *S. cerevisiae* strain BY4741 containing the *dif*-KanMX-*dif* cassette was transformed with yeast vector pAG413GPD (56) expressing wildtype or mutant Lpg1867, or empty vector. Percent excision was quantified by determining the proportion of the transformants that had lost geneticin resistance by plating on selective and nonselective plates.

### Fluorescence microscopy

Overnight cultures of Lp02, Lp02*Δdif*, Lp02*Δlpg1867*, and Lp02*Δlpg1867ΔrecA* were grown from a 2-day old patch in AYET medium. Bacteria were collected at post-exponential phase (OD_600_ 4.0 to 4.5). Approximately ∼1 × 10^9^ cells (1 ODU) from each strain were washed once with 1 mL of 1× PBS before a final resuspension in 500 μL of 1× PBS. The bacteria were stained with 15 μg/mL of DAPI (Roche) for 30 min at room temperature and imaged using a 63× oil immersion lens on a Zeiss Axio Imager.M2 microscope.

### Colony size assay

The distribution of colony size was determined by resuspending 2-day-old patches of the indicated strains in AYET. The OD_600_ of each cell suspension was determined, and the appropriate dilutions made so that ∼200-300 cells were added to each plate. Spread plates were incubated at 37℃ for 4 days before imaging using a SynGene system with GeneSnap software (Syngene). Colony diameter was determined using ColTapp automated image analysis software (57), and plotted using GraphPad Prism version 9.2.0.

### Bacterial infection of U937 macrophages

TPA-differentiated U937 cells were seeded in a 24-well plate at 4 × 10^5^ cells per well in 500 μL of RPMI supplemented with glutamine and 10% heat-inactivated fetal bovine serum. Cells were incubated overnight at 37℃ with 5% CO_2_. Overnight cultures of *Legionella* were grown to post-exponential phase (OD 4.0–4.5 and motile) and used to inoculate the U937 cells at an MOI of 0.05. After 2 h, cells were washed three times with media to remove extracellular bacteria before incubating in fresh media at 37℃ with 5% CO_2_ for a further two days. At 2 h and 48 h post-infection, cells were lysed with 0.02% saponin and then plated on CYET to determine the number of bacterial colony-forming units.

### Bioinformatic analysis of XerL

The amino acid sequence of XerL from *L. pneumophila* was used to uncover putative orthologs of XerL. XerL was queried against the NCBI Legionellales database (taxid:118969) using both BLASTp and tBLASTn with the default settings (58), in addition to a tBLASTn search of additional *Legionella* species that were deposited in the NCBI Sequence Read Archive (59, 60). The resulting putative XerL orthologs can be found in Table S3.

An unrooted phylogeny of Xer proteins was generated using the amino acid sequences from representative XerL orthologs, in addition to reference Xer sequences (Table S3). The amino acid sequences were aligned using MUSCLE (61) and the tree was generated with the FastTree (v 2.1.11) plugin on Geneious Prime using the default settings (62). The resulting tree was visualized using the Interactive Tree of Life (iTOL) server (v 6) (63). A rooted phylogeny of all putative XerL orthologs was generated as described above, using XerD from *Escherichia coli* as the root.

### Materials and Data Availability

All strains and plasmids are available upon request. Sequences and locations of *dif* sites are listed in figures and Tables S1 and S2. The accession numbers and amino acid sequences for Xer proteins are listed in Table S3.

## ACKNOWLEDGEMENTS

We thank members of the Ensminger laboratory for their suggestions and careful reading of the manuscript, and Jordan Lin and John MacPherson for their help with the recombination assays and HHPred analysis. We also thank Hayley Newton for the kind gift of genomic DNA from *C. burnetii* strain nine mile phase II. SRD is supported by a fellowship from the Department of Biochemistry, University of Toronto and an Ontario Graduate Scholarship. This work was supported by a Project Grant from the Canadian Institutes of Health Research (PHT-148819).

## FIGURE LEGENDS

**Figure S1. Phylogenetic analysis of XerL-like recombinases indicates a widespread distribution in Legionellales.** XerL from *L. pneumophila* was used as a query to search for putative XerL orthologs using BLASTp and tBLASTn. The amino acid sequences of these putative orthologs were aligned with MUSCLE. A phylogeny was generated from the alignment using FastTree, with XerD from *E. coli* used as the root. Xer sequences that were also present in Figure 7 are denoted with colored labels as indicated. The scale bar indicates the number of amino acid substitutions per site.

**Table S1. The att site is close to the cumulative GC-skew maximum in all sequenced L. pneumophila strains and Legionella species.** The nucleotide positions of the att site and cumulative GC-skew maximum (cGC skew max) are shown for each L. pneumophila strain and Legionella species with a completed genome sequence available in Genbank. The cGC skew maximum corresponds to the site of replication termination (34). The genes immediately upstream and downstream of each att site are also indicated.

**Table S2. The putative dif site is close to the cumulative GC-skew maximum in all sequenced C. burnetii strains.** The nucleotide positions of the dif site and cumulative GC-skew maximum are shown for each C. burnetii strain with a completed genome sequence available in Genbank.

**Table S3. The accession numbers and amino acid sequences for the Xer proteins used in the phylogenetic analyses.**

