## Supplementary figures and images for "Mobile element integration reveals a chromosome dimer resolution system in Legionellales"

### Figure S1

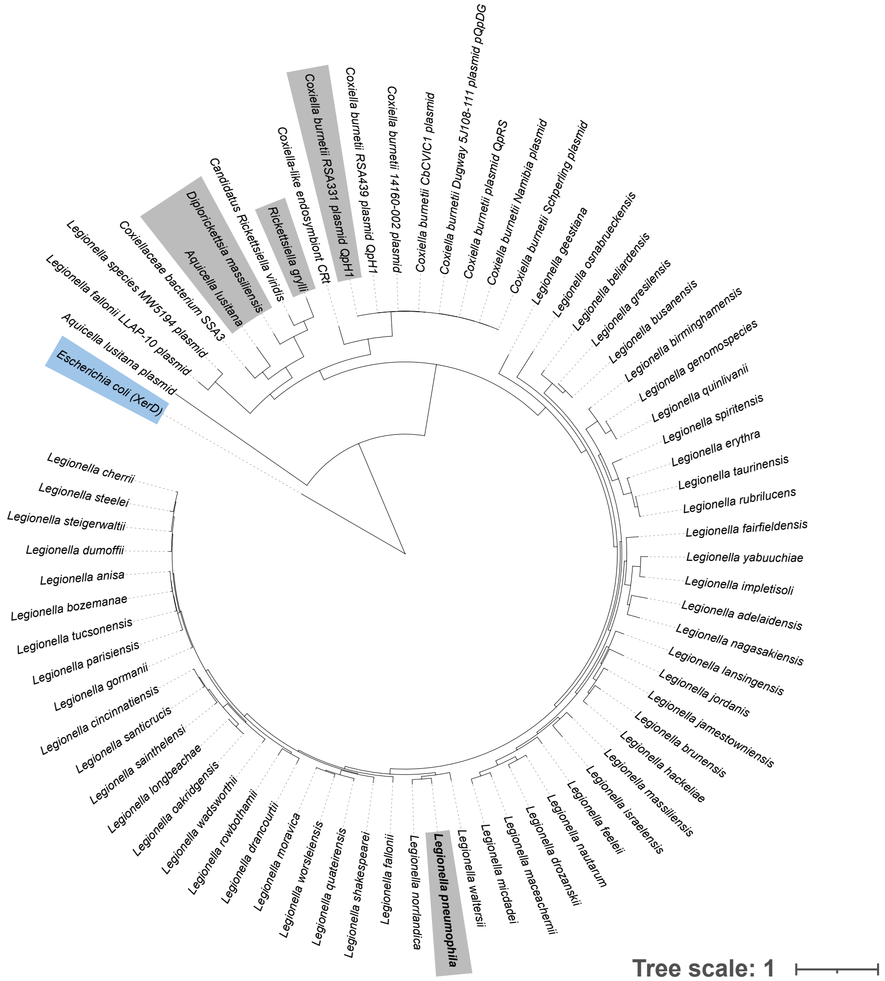
